# TGR5-mediated Ca^2+^ signaling in cholangiocytes

**DOI:** 10.1101/2025.01.23.634480

**Authors:** Xuanmeng Chen, Amr Al-Shebel, Thibault Pebrier, Thierry Tordjmann, Olivier Dellis

## Abstract

The Bile Acid TGR5 receptor is well known to active the cAMP pathways leading to CFTR activation and Cl^−^ ions secretion, needed for bile alkalinization and hydration. However, during cystic fibrosis development, only 10 to 15% of the patients present liver defect due to bile duct disorders, meaning that another process should compensate for the loss of CFTR activity. Interestingly, TGR5 stimulation has also been reported to mobilize Ca^2+^ ions. Using normal human cholangiocytes and cholangiocarcinoma cell lines, we confirmed by using a specific agonist, that TGR5 stimulation induced a Ca^2+^ release from the endoplasmic reticulum and an influx of extracellular Ca^2+^ ions. Next, this Ca^2+^ mobilization allows an ATP (and UTP) release, leading to the activation of P2Y receptors, reinforcing this Ca^2+^ mobilization. This study shows that activation of the BA receptor TGR5 has the capacity to induce the two main intracellular pathways, cAMP and IP_3_-Ca^2+^ in cholangiocytes. From our data, we speculate that the pathway we described will allow activation of the Ca^2+^-activated Cl^−^ channels TMEM16A, in parallel to CFTR in non-CF cells, or to compensate in part or in totality the loss of CFTR in CF patients.

**HIGHLIGHTS:**

- Bile acid receptor TGR5 induces Ca^2+^ mobilization in cholangiocytes
- Ca^2+^ ions come from the endoplasmic reticulum and from the extracellular medium
- P2Y receptors are trans-activated by TGR5 and reinforce the Ca^2+^ mobilization
- This Ca^2+^ pathways might compensate the CFTR defect in CF patients

## 1. INTRODUCTION

Cholangiocytes are biliary epithelial cells lining the biliary tree i.e the intrahepatic and extrahepatic bile ducts as well as the gallbladder. Even if these cells represent ≍ 5 % of the total liver cells, they play a crucial role in bile formation thanks to the expression of several kinds of membrane transporters, ion exchangers and surface receptors, coupled to diverse signaling pathways (review in [1]). For example, secretin receptor stimulation induces cyclic-AMP (cAMP) production by adenylyl cyclase [2, 3] that activates protein kinase A (PKA) which phosphorylates CFTR channels and drives Cl^−^ ions secretion in bile [4], coupled with AE2 (Anion Exchanger 2)-dependent HCO_3_^−^ ions secretion [5]. In combination with this net bicarbonate secretion, cAMP-dependent aquaporin opening results in an increase in ductular bile flow [6] [7]. Of note, in a significant proportion of cystic fibrosis (CF) patients, CFTR defect impairs ductular bile secretion leading to ductal cholestasis ultimately ending in sclerosing cholangitis and cirrhosis [8]

Beside cAMP, intracellular Ca^2+^ ions can also act as second messengers regulating ductular secretion [1]. In particular, cholangiocytes express a large spectrum of ATP purinergic receptors, from ATP-gated cationic channels (P2X receptors) to G protein-coupled receptors (GPCR) (P2Y receptors), the stimulation of which increases cytosolic Ca^2+^ ions concentration ([Ca^2+^]_cyt_) [9]. As ATP is found in bile at micromolar concentrations [10, 11], it can induce in cholangiocytes a Ca^2+^ entry and/or an IP_3_-mediated Ca^2+^ release from the endoplasmic reticulum (ER), respectively through P2X or P2Y receptors. Recently, it has been shown that Cl^−^ ions can pass through TMEM16A (also known as ANO1) which is functionally activated by an increase in ([Ca^2+^]_cyt_ [12, 13]. Therefore, beside the well-known cAMP signaling pathway, Ca^2+^ mobilization should also be crucial for cholangiocyte physiology and pathophysiology. Even if bile acids (BAs) where known to induce Ca^2+^ mobilization in cholangiocytes [14], it has been mostly reported before the discovery of their membrane receptor TGR5 [15] (see below). Interestingly, even if the IP_3_/Ca^2+^ pathway has been clearly characterized after P2Y stimulation with ATP in cholangiocytes [9], nothing is known about a putative BA-induced TGR5-mediated Ca^2+^ signaling pathway in those cells.

The BA receptor TGR5, also known as GPBAR1 (G Protein-coupled BA Receptor-1) is the most studied membrane BA receptor [16]. High levels of TGR5 mRNA were detected in organs like gallbladder, intestine, stomach and spleen (review in [17]). In the liver, TGR5 is poorly expressed in hepatocytes, but highly in cholangiocytes [16, 17]. TGR5 has been reported to be involved in the regulation of metabolism [18, 19], inflammation [19], liver regeneration [20, 21] and cancer cell proliferation [17].

TGR5 activation is mostly reported to drive cAMP production [22], leading to downstream activation of diverse signaling pathways like NF-κB, AKT, CEBPb and Erk [17], with impact on cell proliferation [23], migration, and apoptosis in cholangiocytes (review in [24]. TGR5 is also reported to be involved in CFTR-dependent Cl^−^ ions secretion [16]. Noteworthy, TGR5 expression is increased in cholangiocarcinomas where it could favor cholangiocyte proliferation and decrease apoptosis [24–26], whereas it is less expressed in Primary Sclerosing Cholangitis (PSC) patient liver where this reduction contributes to aggravate biliary injury [27].

In this work, we studied intracellular Ca^2+^ homeostasis in normal cholangiocytes and in commonly used cholangiocarcinoma cell lines under stimulation with a specific TGR5 agonist developed by Roche [28], which belongs to the 1-hydroxyamino-3-phenyl-propanes family. Our results clearly show that specific TGR5 stimulation induces a Ca^2+^ release from internal stores and a subsequent Ca^2+^ influx through complex and multiple pathways involving Gαq/11, P2X and P2Y receptors, and a Store-Operated Calcium Entry (SOCE). Our study pointed out that beside the well reported cAMP pathway, BA signaling can be transduced through TGR5-mediated Ca^2+^ responses, with potential impact on bile physiology and pathophysiology. We also unveiled differences in TGR5-induced Ca^2+^ mobilization between normal and malignant cholangiocytes. Altogether this work may provide interesting targets for further pathophysiological investigation and future therapeutic interventions.

## 2. MATERIALS & METHODS

### 2.1. Cell lines

Normal human cholangiocytes were from Cliniscience and maintained in a special and complete medium from Cliniscience (Nanterre, France).

Human intrahepatic HuCCT1 cell line was a gift from Dr Cedric Coulouarn (INSERM U1241, Rennes, France). Human extrahepatic Mz-Cha1 cell line was a gift from Dr Laura Fouassier (Centre de Recherche Saint Antoine, INSERM U938, Paris, France). Rat intrahepatic F258 cell line was a gift from Dr Dominique Lagadic-Gossman (Irset, Rennes, France). HuCCT1 cells lines were basically maintained in RPMI-1640 + Glutamax medium, Mz-Cha1 in DMEM + Glutamax medium and F258 in William’s medium. All our media were from Lonza (Verviers, Belgium) and supplemented with 10 % heat-inactivated fetal calf serum. F258 cell medium culture was also supplemented with 2 mM L-Glutamine (Lonza, Verviers, Belgium), Mz-Cha1’s with 1% Hepes.

Normal human cholangiocytes and cholangiocarcinoma cell lines were maintained at 37°C in a 5% CO_2_ –humidified atmosphere.

### 2.2. Cytosolic Ca^2+^ concentration

[Ca^2+^]_cyt_ was recorded by a fluorimetric ratio technique as in [29, 30]. Culture medium was removed, cells were washed with PBS and Trypsin-EDTA was added 5-10 min to detach the cells from the support. Next, fresh culture medium was added to stop the trypsin action, and the cells were spun. After removal of the supernatant, the cells were resuspended at a density of 10^6^ cells/ml in Ca^2+^-free Hepes Buffered Saline solution (HBS; 135 mM NaCl, 5.9 mM KCl, 1.2 mM MgCl_2_, 11.6 mM Hepes, 11.5 mM glucose adjusted to pH 7.3 with NaOH) and incubated in the dark with 4 μM Indo-1-AM (cholangiocarcinoma cell lines) or Fura-2-AM (Normal human cholangiocytes) for 45 min at room temperature under slow agitation. Cells were then centrifuged and resuspended in dye-free HBS medium supplemented with 1 mM CaCl_2_. For experiments, 0.5 to 1 × 10^6^ cells were suspended in 2 ml HBS in a quartz cuvette and inserted into a spectrofluorophotometer (RF-1501 Shimadzu Corporation, Kyoto, Japan) connected to a PC computer (Dell Computer Corp., Montpellier, France). A temperature of 37°C was maintained by circulating water from a thermostatic bath. For the cholangiocarcinoma cell lines, ultraviolet light of 360 nm was used for excitation of Indo-1, and emissions at 405 and 480 nm were recorded. For the normal human cholangiocytes, Fura-2 was excited at 340 and 380 nm, and emission at 510 nm was recorded. Background and autofluorescence of the cell suspension were subtracted from the recordings. The maximum dye fluorescence (*R*_max_) was obtained by adding 1 μM ionomycin to the cell suspension in the presence of 10 mM CaCl_2_. Minimum fluorescence was determined following depletion of external Ca^2+^ with 5 mM EGTA. [Ca^2+^]_cyt_ was calculated according to the equation [Ca^2+^]_cyt_ = *K*_d_ (R-*R*_min_)/(*R*_max_-R)F, where *K*_d_ is the apparent dissociation constant of the Ca^2+^-dye (230 nM for Indo-1, 250 for Fura-2), R is the ratio of fluorescence at 405 and 480 nm for Indo-1 (ratio of fluorescence emitted at 510 nm for Fura-2) and F is the ratio of fluorescence values recorded at 480 nm in absence and presence of 10 mM CaCl_2_ (at 510 nm for Fura-2 after excitation at 380 nm) [29].

Some experiments were conducted in absence of external Ca^2+^ ions and in presence of 1 mM EGTA to study the role of the internal Ca^2+^ stores in the cell Ca^2+^ homeostasis. The other experiments were conducted in presence of 1 mM external CaCl_2_.

### 2.3. Chemicals

All our chemicals were from Sigma (Saint Quentin Fallavier, France). RO5527239 was a kindly gift of Roche [28], and it specificity for TGR5 was previously determined by our laboratory [21, 26].

### 2.4. Statistical analysis

Given values are representative of at least 3 independent experiments and are given as mean ± SEM. When used, a t-test < 0.05 is considered as significant. In pharmacological experiments, a paired Student test was used, and a value < 0.05 was considered as significant.

## 3. RESULTS

### 3.1. TGR5-specific stimulation induces intracellular Ca^2+^ signals in normal human cholangiocytes and cholangiocarcinoma cell lines

We characterized Ca^2+^ homeostasis in normal human cholangiocytes (NHC), and in two commonly used human cholangiocarcinoma (CCA) cell lines, one extrahepatic (Mz-Cha1) and one intrahepatic (HuCCT1). We also used the rat liver biliary epithelial cell line F258 [31]. First, we found that in the presence of 1 mM extracellular Ca^2+^ ions, the resting cytosolic Ca^2+^ concentration (rest[Ca^2+^]_yt_) was significantly smaller (p < 0.05, n > 50) in the 3 cell lines (149 ± 4, 117 ± 3 and 150 ± 5 nM), respectively for Mz-Cha1, HuCCT1 and F258, than in NHC (184 ± 11 nM).

Second, we found that the TGR5-specific agonist RO5527239 can mobilize Ca^2+^ ions in NHC and in the 3 cell lines in our conditions. As shown in figure 1A, TGR5-dependent Ca^2+^ mobilisation was transient, and increasing RO5527239 concentrations induced progressively higher Ca^2+^ mobilisation, with a plateau for concentrations > 90-120 µM for the 3 cell lines, *vs*. > 300 µM in NHC (Fig. 1A&B). Dose-response curves fitting gave an *EC*_50_ ≈ 65 µM in the 3 cell lines (71.4 ± 3.2, 66.1 ± 2.4 and 62.8 ± 3.6 µM respectively for Mz-Cha1, HuCCT1 and F258 cells, Fig. 1B), whereas the *EC*_50_ for NHC was significantly higher (127.1 ± 10.1 µM, n =3, p < 0.01), twice more than for CCA cell lines (Fig. 1B). This difference likely reflects the fact that TGR5 expression is increased in CCA as compared with normal cholangiocytes [23, 32]. Noteworthy, despite the fact that higher RO5527239 concentrations were needed to rise the maximum of Ca^2+^ mobilisation in NHC, the Ca^2+^ mobilisation amplitude in NHC cells was largely higher than in CCA cell lines: 567 ± 47 nM for NHC *vs*. 277 ± 19, 237 ± 23 and 161 ± 28 nM respectively for Mz-Cha1, HuCCT1 and F258 cells (n > 20, Fig. 1C).

**Figure 1:**
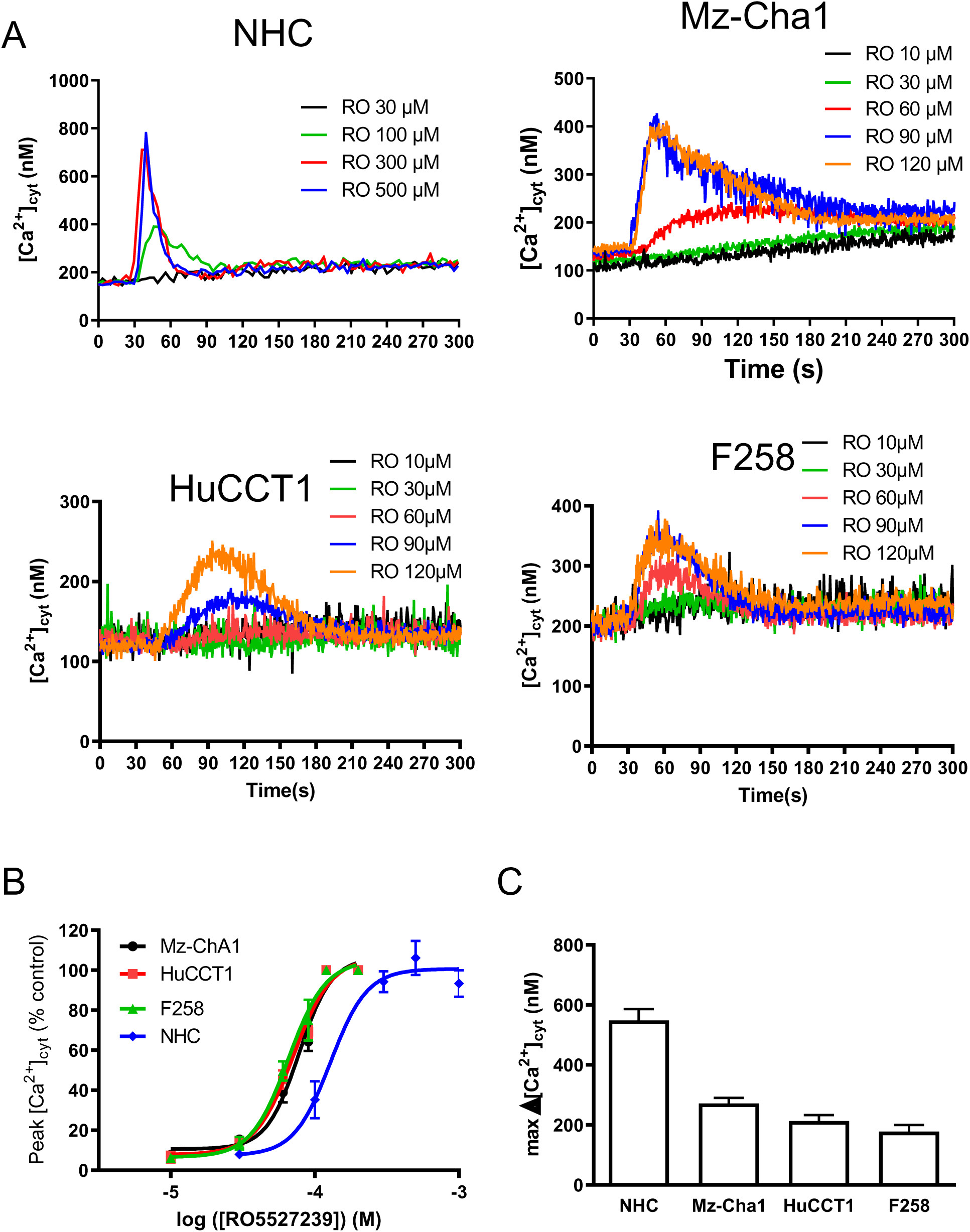
The TGR5 specific agonist RO5527239 induces calcium mobilization of normal human cholangiocytes, and cholangiocytes cell lines. A. Cytosolic Ca^2+^ concentration ([Ca^2+^]_cyt_) measurement of Normal Human Cholangiocytes (NHC), Mz-Cha1, HuCCT1 and F258 cell lines using Fura-2 or Indo-1 fluorescence. Cells were treated with increasing concentration of RO5527239 (“RO”) added at t = 30 s. B. RO5527239 dose-response curves could be fitted with a simple sigmoidal curve, giving an *EC*_50_ of 71.4 ± 3.2 µM (Mz-Cha1 cells), 66.1 ± 2.4 µM (HuCCT1 cells), 62.8 ± 3.6 µM (F258 cells) and 127.1 ± 10.1 µM (NHC), n =3 for each. C. Maximal cytosolic calcium concentration variation (Δ[Ca^2+^]_cyt_) induced by optimal RO5527239 concentration (120 µM for the cell lines, 300 µM for NHC cells).

To clearly establish that RO5527239-mediated Ca^2+^ mobilisation was TGR5-specific, we performed experiments with JTE-013, a specific inhibitor of S1PR, which is known to bind BAs [33]. In our conditions, JTE-013 was unable to reduce this Ca^2+^ mobilisation, showing that BAs induced Ca^2+^ through a S1PR2-independent pathways (not shown).

Altogether these observations suggest that BA-mediated TGR5 activation might directly or indirectly operate a control on intracellular Ca^2+^ homeostasis in cholangiocytes.

### 3.2. TGR5 stimulation induces a Ca^2+^ release and a Ca^2+^ influx

To more deeply understand mechanisms involved in TGR5-mediated Ca^2+^ responses, we first removed extracellular Ca^2+^ ions before TGR5 stimulation. As clearly shown in figure 2A, the absence of extracellular Ca^2+^ ions almost suppressed the Ca^2+^ mobilisation in NHC and Mz-Cha1 cells (not shown for HuCCT1 and F258 cells). In line, Ca^2+^ peaks and areas under curves (AUC) upon TGR5 stimulation were decreased by ≈ 80% in the lack of extracellular Ca^2+^ as compared with 1 mM external Ca^2+^ (−80.5 ± 3.8, −86.5 ± 3.0, −71.6 ± 4.5 and −77.9 ± 1.5 % respectively for NHC, Mz-Cha1, HuCCT1 and F258 cells, n >3, Fig. 2B). These results thus showed that TGR5 stimulation not only induced a massive Ca^2+^ entry, but also a Ca^2+^ release from internal stores. Although this Ca^2+^ release looked like quantitatively small, data from Figure 2C indicated that it comes from the endoplasmic reticulum (ER). As shown in Figure 2C, we pre-treated the NHC and Mz-Cha1 cells with thapsigargin (TG), an irreversible ER Ca^2+^ (SERCA) pump inhibitor known to induce a passive Ca^2+^ release from the ER, and a subsequent Ca^2+^ influx known as a “Store-Operated Calcium Entry” (SOCE [34]) if extracellular Ca^2+^ ions are present. In the absence of extracellular Ca^2+^ (Fig. 2C, black curves), the SOCE is abolished and the [Ca^2+^]_cyt_ increase is only due to Ca^2+^ release from the ER (ER Ca^2+^ emptying). Importantly, subsequent stimulation with RO5527239 (“RO”, 90 µM for the cell lines, 300 µM for NHC cells) did not elicit any additional [Ca^2+^]_cyt_ increase, showing that after ER Ca^2+^ depletion, TGR5 stimulation cannot further induce any Ca^2+^ signal. Ca^2+^ release from the ER thus appears as a crucial step in the TGR5-mediated Ca^2+^ signalling pathway. In line with these data, in cells pre-treated with TG, in the presence of extracellular Ca^2+^ ions (Fig. 2C, red curves), RO5527239 was neither able to increase [Ca^2+^]_cyt_; interestingly, RO5527239 rather induced a [Ca^2+^]_cyt_ decrease, possibly due to an activation of Ca^2+^ efflux from the cells through PMCA/NCX [35]. The same results were obtained on HuCCT1 and F258 cells (not shown).

**Figure 2:**
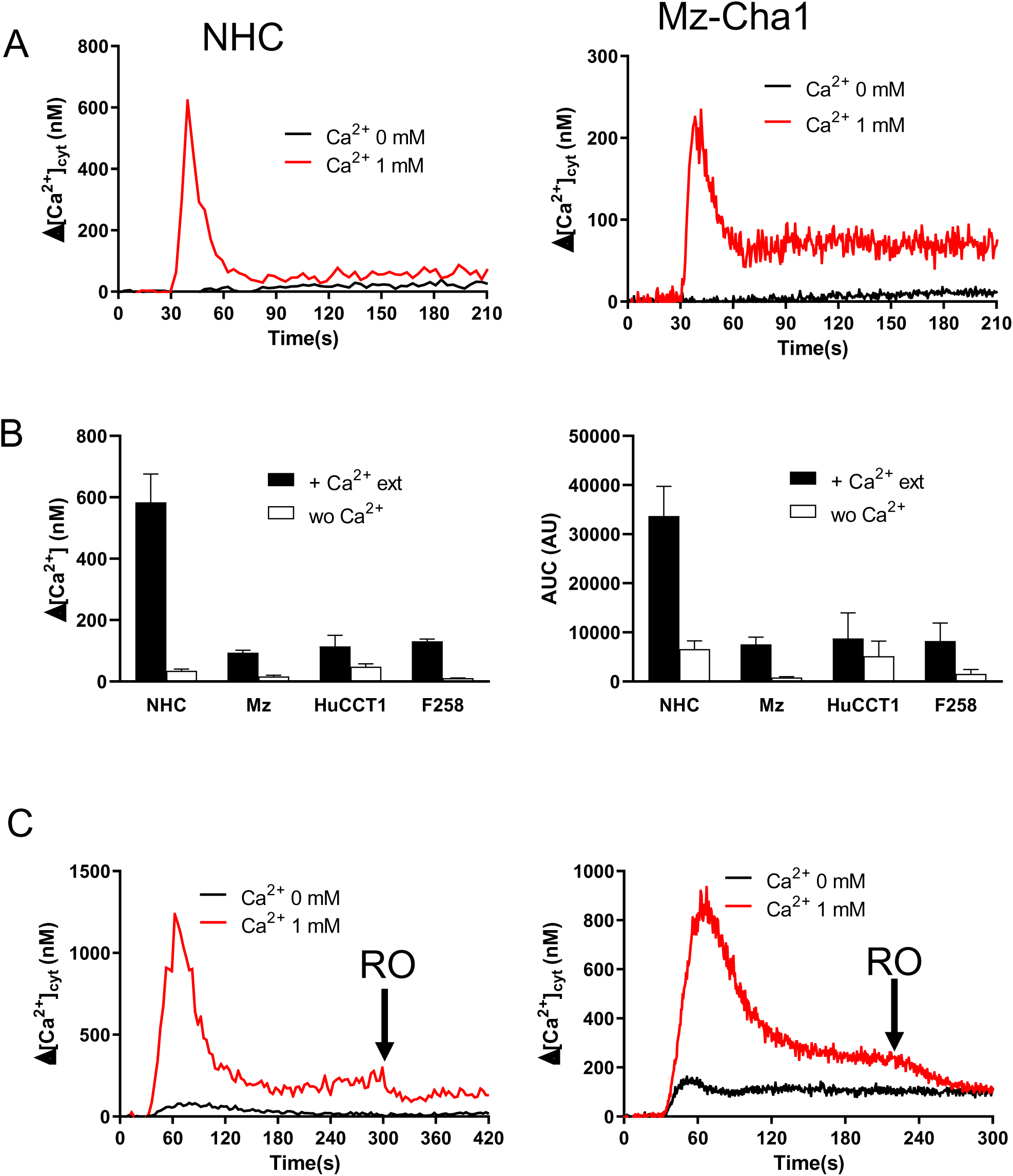
TGR5 stimulation mobilizes intracellular calcium and external calcium. A. Cytosolic Calcium concentration variation (Δ[Ca^2+^]_cyt_) of NHC and Mz-Cha1 cells, in presence (red curves) or absence (black curves) of external 1 mM CaCl_2_, after stimulation by 300 or 90 µM RO5527239 respectively in NHC and Mz-Cha1 cells added at t = 30 s. B. Maximal cytosolic calcium concentration variation (Δ[Ca^2+^]_cyt_, left) and area under the curve (AUC, right) induced by optimal RO5527239 concentration (90 µM for the cell lines, 300 µM for NHC cells) in absence “wo Ca^2+^”) or presence (“+ Ca^2+^ ext”)of external 1 mM CaCl_2_ (n > 3) C. Δ[Ca^2+^]_cyt_ of NHC and Mz-Cha1 cells treated by 1 µM thapsigargin at t=30s in absence or presence of external 1 mM CaCl_2_. At t=300s, 90 µM or 300 µM RO5527239 was added respectively for Mz-Cha1 cells and NHC (black arrow)

### 3.3. TGR5 stimulation activates a Gαq/11 pathway

As Ca^2+^ release from the ER appeared to play a major role in TGR5-mediated Ca^2+^ signalling, further exploration of the involved transduction pathway was performed. To explore the IP_3_ pathway, we pre-treated the cells with 2 inhibitors: U73122, a well-known phospholipases (PL) inhibitor impairing IP_3_ synthesis, and YM254890, a new Gαq/11 inhibitor impairing PL activation. Both inhibitors inhibited 90-95% of the TGR5-mediated Ca^2+^ signal (Fig. 3). These results clearly established that TGR5 stimulation activates a Gαq/11 pathway, leading to PL activation and IP_3_ synthesis, allowing Ca^2+^ release from the ER and a subsequent SOCE.

**Figure 3:**
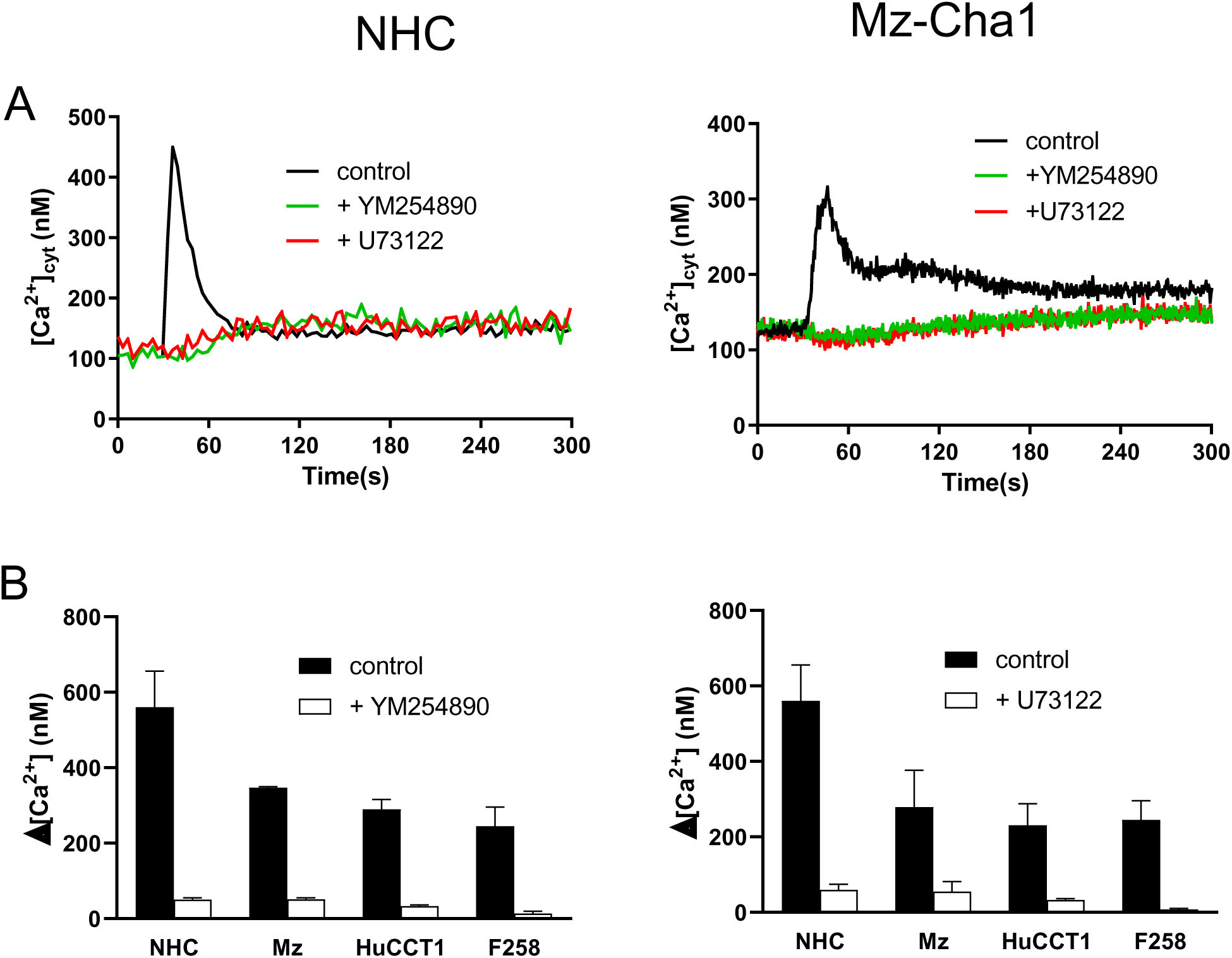
TGR5-induced calcium mobilization is phospholipase dependant. A. Cytosolic Calcium concentration ([Ca^2+^]_cyt_) of NHC (left) and Mz-Cha1 (right) cells stimulated by 300 or 90 µM RO5527239 respectively, in control conditions (black curves), or after a 10 min - pretreatment by the specific inhibitor of Gαq/11 YM254890 (1 µM, green curves), or after a 10 min – pretreatment by the specific inhibitor of phospholipases U73122 (5 µM, red curves). B. Maximal cytosolic calcium concentration variation (Δ[Ca^2+^]_cyt_) induced by optimal RO5527239 concentration (90 µM for the cell lines, 300 µM for NHC cells) in control conditions or after pretreatment by YM254890 (1 µM, left) or U73122 (5 µM, right).

### 3.4. TGR5-induced Ca^2+^ mobilization is modulated by P2Y receptors

Recently it has been shown that millimolar concentrations of the two BA Ursodeoxycholic acid (UDCA) and Tauro-UDCA (TUDCA) induce a [Ca^2+^]_cyt_ rise in cholangiocytes which is dependent on IP_3_ receptors and ATP-mediated P2 receptors [13]. We thus hypothesized that TGR5 stimulation might elicit ATP release and added apyrase before RO5527239 challenge. We observed that TGR5-mediated Ca^2+^ response was partly blunted after apyrase treatment (Fig. 4A), confirming that extracellular NTP play a role in TGR5-induced Ca^2+^ signalling. Noteworthy, bile contains adenosine nucleotides, and the Mz-Cha1 cell line has been reported to secrete ATP [10]. Furthermore, when secreted, ATP is accompanied by UTP in a ratio of 3 for 1 [36], meaning that in bile, not only ATP can be active, but also UTP.

**Figure 4:**
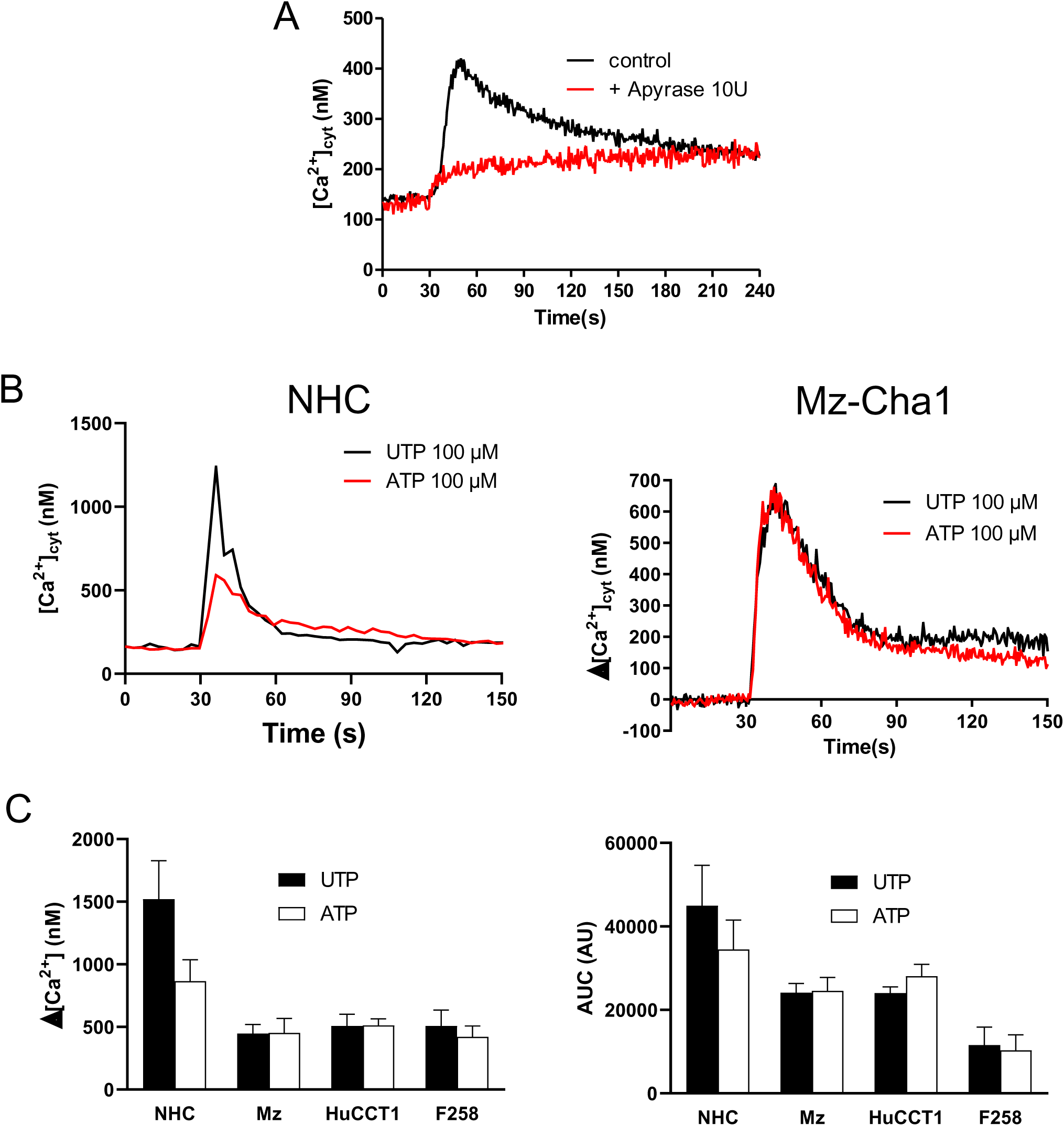
ATP (UTP) and P2Y2R receptors are implied in TG5-induced calcium mobilization. A. Cytosolic Calcium concentration variation (Δ[Ca^2+^]_cyt_) of Mz-Cha1 cells stimulated by 90 µM RO5527239 in presence of apyrase 10 U. B. Δ[Ca^2+^]_cyt_ of NHC and Mz-Cha1 cells treated by 100 µM UTP (black curves) or ATP (red curves) at t=30s. C. Maximal cytosolic calcium concentration variation (Δ[Ca^2+^]_cyt_, left) and area under the curve (AUC, right) induced by optimal RO5527239 concentration (90 µM for the cell lines, 300 µM for NHC cells) in absence or presence of external 1 mM CaCl_2_ (n > 3)

As cholangiocytes express various types of P2X and P2Y receptors [37–41], we next tested P2 receptors inhibitors on TGR5-mediated Ca^2+^ responses. Interestingly, if cholangiocytes express several P2Y receptors transcripts, only P2Y2 receptor (P2Y2R) protein is reported to be expressed in these cells [38]. As shown in figure 4B, 100 µM ATP and 100 µM UTP gave the same Ca^2+^ signals in the Mz-Cha1 cell line, with similar kinetics (Ca^2+^ peak and area under curves). As only P2Y2R receptor is equally activated by ATP and UTP (reviewed in [42]), our data suggest that P2Y2R was probably responsible for ATP (and UTP)-induced Ca^2+^ signals in our cell lines (the same results were obtained with HuCCT1 and F258 cells, not shown). Interestingly, the Ca^2+^peak amplitude was almost 1.7-fold higher with UTP than with ATP in NHC cells (1521 ± 307 *vs.* 865 ± 170 nM, n =3), suggesting that UTP could activate another UTP-sensitive P2 receptor (like P2Y4R) in these cells, which is not activated by ATP [42]. Experiments performed in the lack of extracellular Ca^2+^ ions (Fig 5) showed that ATP and UTP can induce a Ca^2+^ release and a Ca^2+^ influx, what we expected for a GPCR like P2Y2R (and possibly P2Y4R in NHC).

**Figure 5:**
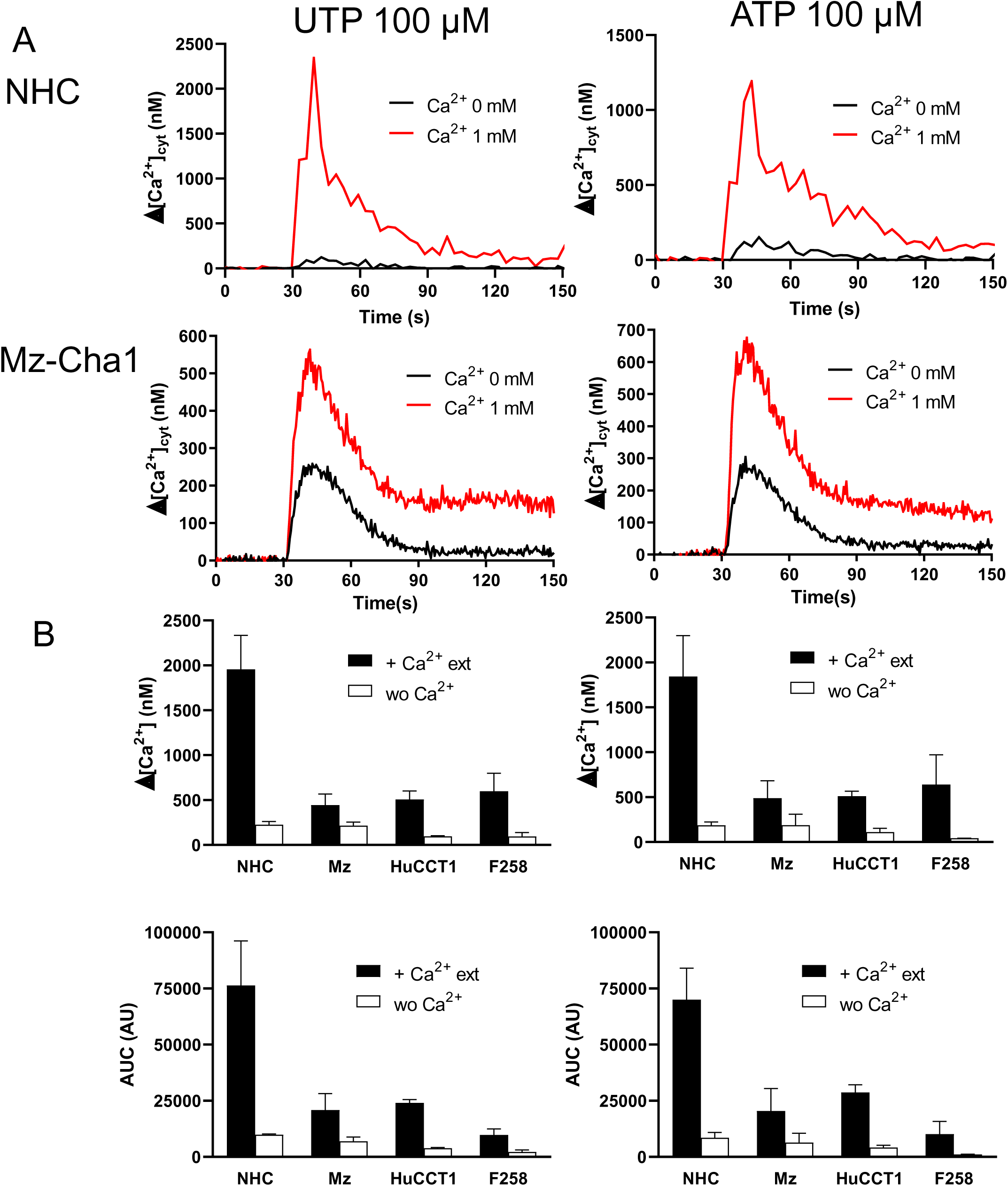
UTP/ATP mobilizes intracellular calcium and external calcium. A. Cytosolic Calcium concentration variation (Δ[Ca^2+^]_cyt_) of NHC (top) and Mz-Cha1 cells (bottom), in presence (red curves) or absence (black curves) of external 1 mM CaCl_2_, after stimulation by 100 µM UTP (left) or ATP (right). B. Peak (top) and area under curves (bottom) of UTP (left) and ATP (right) induced calcium mobilisation in the 4 cell types.

In YM254890-pre-treated cells, UTP was no longer able to induce any Ca^2+^ mobilisation (Fig 6A&B), confirming that ATP and UTP activated P2Y receptors. Noteworthy, ATP-induced calcium signals were totally impaired by Gαq/11 inhibition (Fig. 6B), suggesting that in these cells, ATP cannot induce any detectable Ca^2+^ influx through P2X receptors.

**Figure 6:**
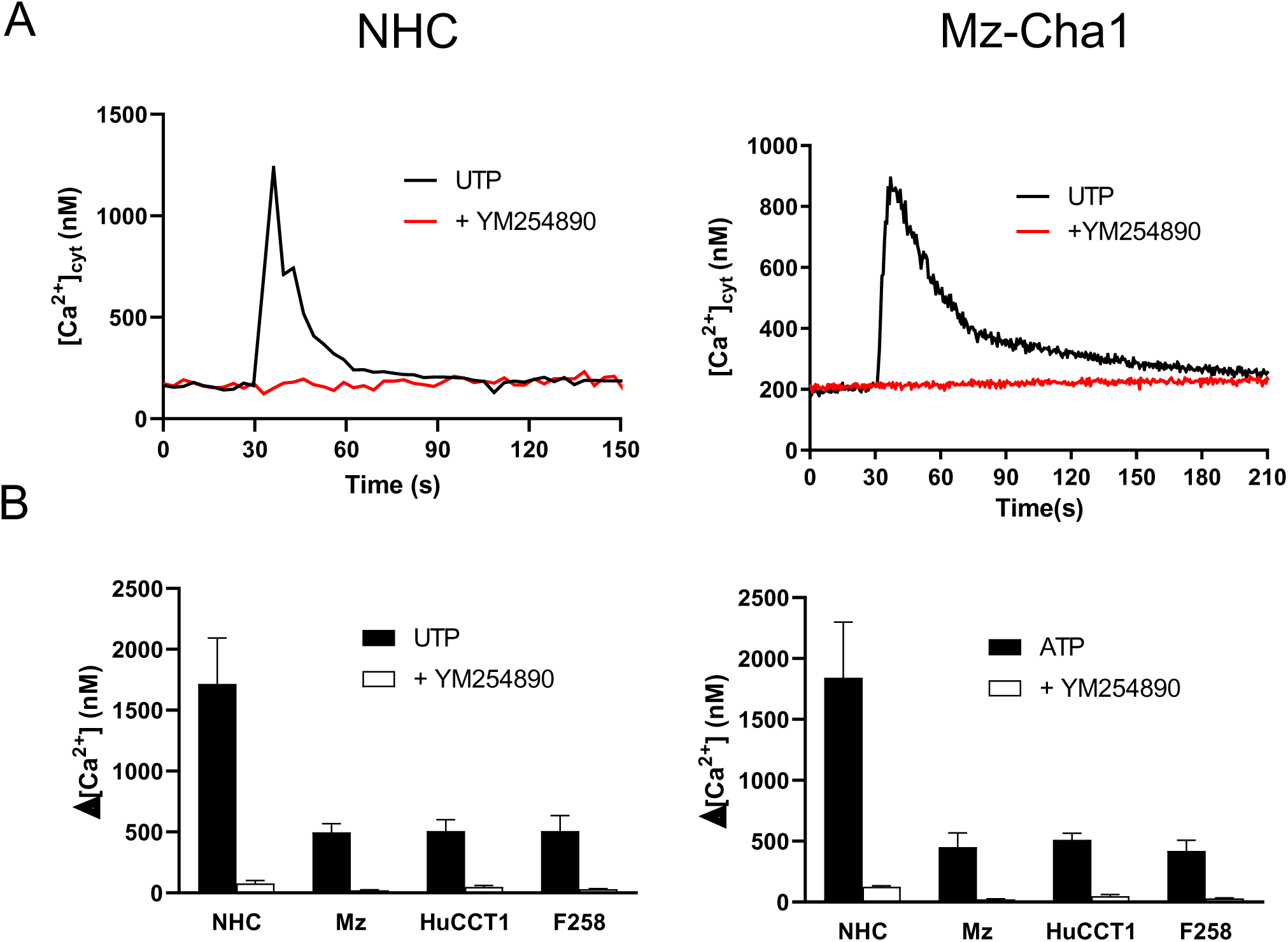
UTP-induced calcium mobilisation is Gαq/11 dependent. A. Cytosolic Calcium concentration variation (Δ[Ca^2+^]_cyt_) of NHC (left) and Mz-Cha1 cells (right), stimulated by UTP (100 µM alone, black curves) or after a 10 min - pretreatment by the Gαq/11 specific inhibitor YM254890 (1 µM, red curves) B. Peak of UTP (left) and ATP (right) induced calcium mobilisation in the 4 cell types in control conditions or after inhibition of Gαq/11.

Next, in the 3 cell lines, the specific P2Y2R inhibitor ARC118925XX induced a >95% inhibition of ATP or UTP-induced Ca^2+^ mobilisation, confirming that in these cells, ATP/UTP induced Ca^2+^ mobilisation through P2Y2R (Fig. 7A&B). However, in NHC, ARC118925XX almost totally abolished ATP-induced Ca^2+^ mobilisation but only decreased by ≍2 thirds UTP-induced Ca^2+^ responses (1715 ± 377 *vs.* 540 ± 129 nM, n = 3). These data suggest that in NHC, ATP induced Ca^2+^ mobilisation through P2Y2R (like in the 3 cell lines), and that UTP induced a second Ca^2+^ mobilisation probably through P2Y4R.

**Figure 7:**
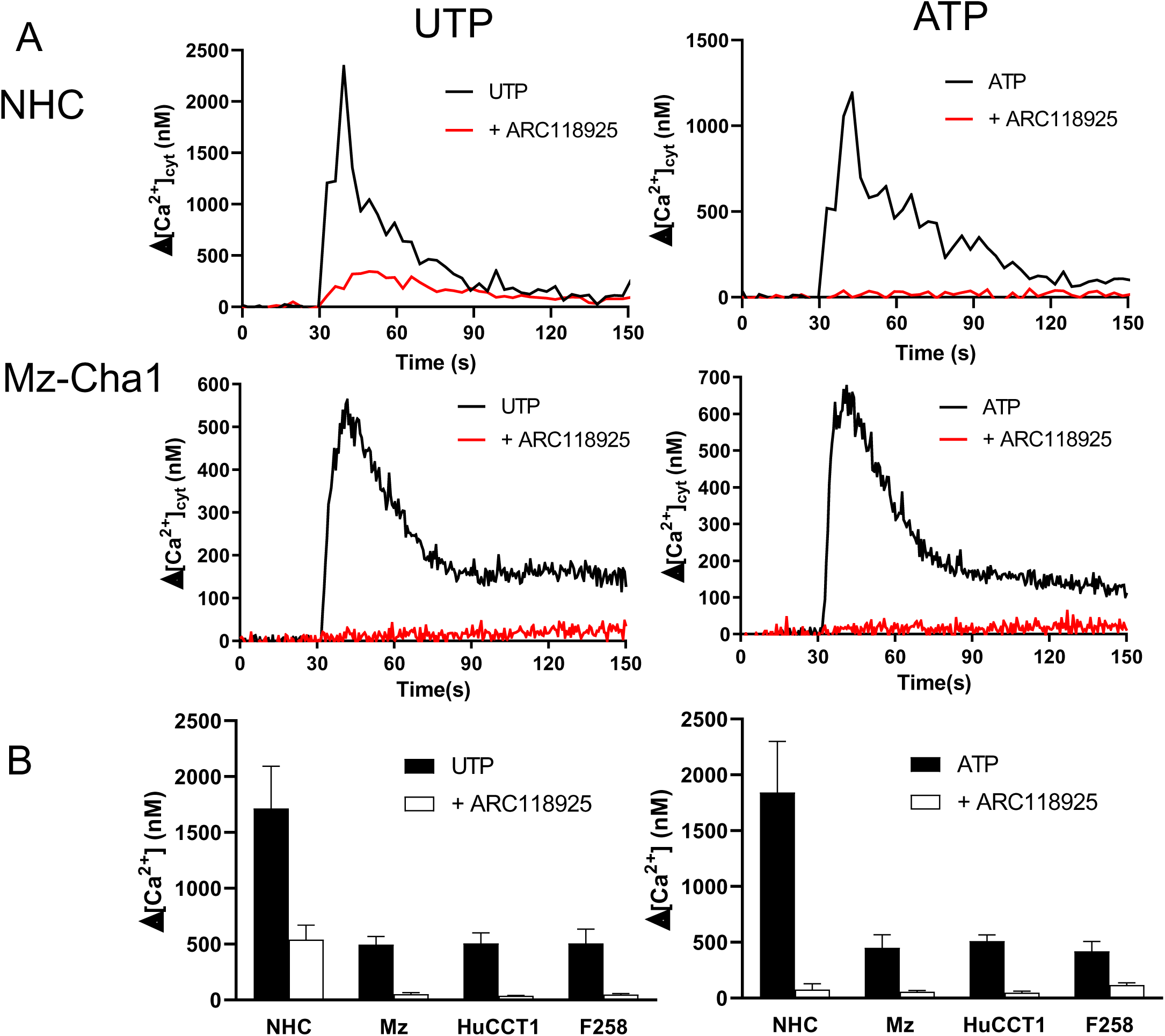
UTP/ATP-induced calcium mobilisation is P2Y2RR (and P2Y4RR?) dependent. A. Cytosolic Calcium concentration variation (Δ[Ca^2+^]_cyt_) of NHC (top) and Mz-Cha1 cells (bottom), stimulated by UTP (100 µM, left) or ATP alone (100 µM, right, black curves) or after a 10 min - pretreatment by the P2Y2RR specific inhibitor ARC118925 (10 µM, red curves) B. Peak of UTP (left) and ATP (right) induced calcium mobilisation in the 4 cell types in control conditions or after inhibition of P2Y2RR.

On these bases, we treated cells with the P2Y2R receptor inhibitor ARC118925XX before TGR5 stimulation and observed that the TGR5-mediated Ca^2+^ peak was decreased by ≍ 50 % (−50.5, −33.2, −49.0 and −56.8 % respectively for NHC, Mz-Cha1, HuCCT1 and F258 cells, Fig. 8A&B), highlighting that TGR5-induced Ca^2+^ response is partly due to P2Y2R receptors involvement.

**Figure 8:**
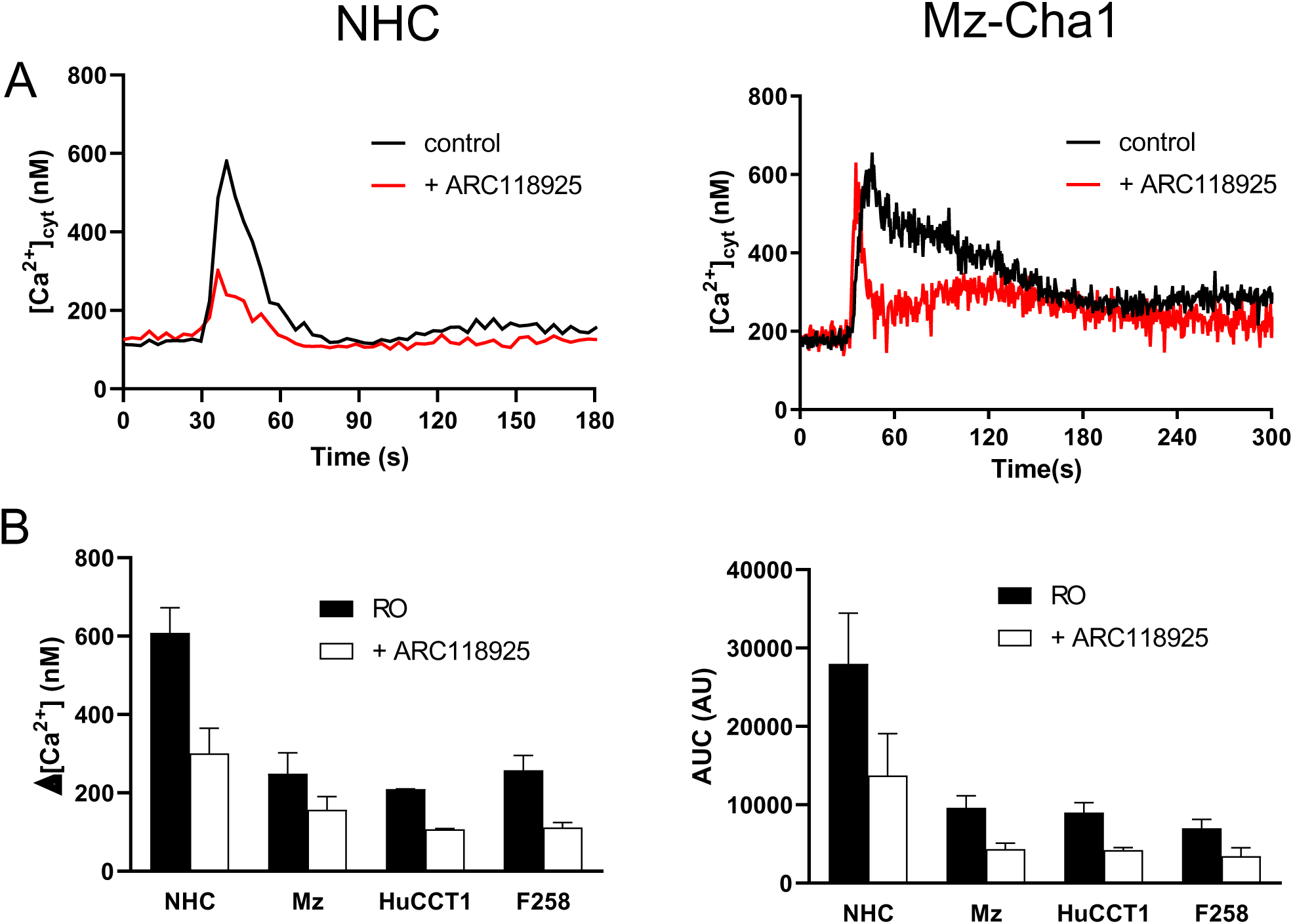
TGR5 – induced calcium mobilization is Gαq/11 and P2Y2RR-dependent. A. Δ[Ca^2+^]_cyt_ of NHC and Mz-Cha1 cells treated by 300 or 90 µM RO5527239 respectively in control conditions (black curves), or after a 10 min – pretreatment by the P2Y2RR inhibitor ARC118925XX (10µM, red curves). B. Peak (left) and area under curves (right) of the RO5527239-induced calcium mobilisation in control conditions or after pretreatment by the specific P2Y2RR inhibitor ARC118925 (10 µM) in the 4 cell types.

Although data presented above suggest that ATP-induced Ca^2+^ responses were exclusively dependent on P2Y2R in our cells, we tested for P2X4R contribution, as this receptor is highly expressed in normal rat cholangiocytes (NRC) and Mz-Cha1 cells, and increases the trans-epithelium conductance after ATP stimulation [43]. Surprisingly, specific P2X4R inhibition with 5-BDBD or BAY1797 potentiated TGR5-induced Ca^2+^ mobilisation by 2.7, 2.1, 1.8 and 1.4-fold respectively for NHC, Mz-Cha1, HuCCT1 and F258 cells (Fig. 9A&B). Thus, P2X4R has a complex role in the TGR5-induced Ca^2+^ mobilisation, not acting as a Ca^2+^ entry pathways (Fig. 10), but more as a regulator of the Ca^2+^ mobilisation.

**Figure 9:**
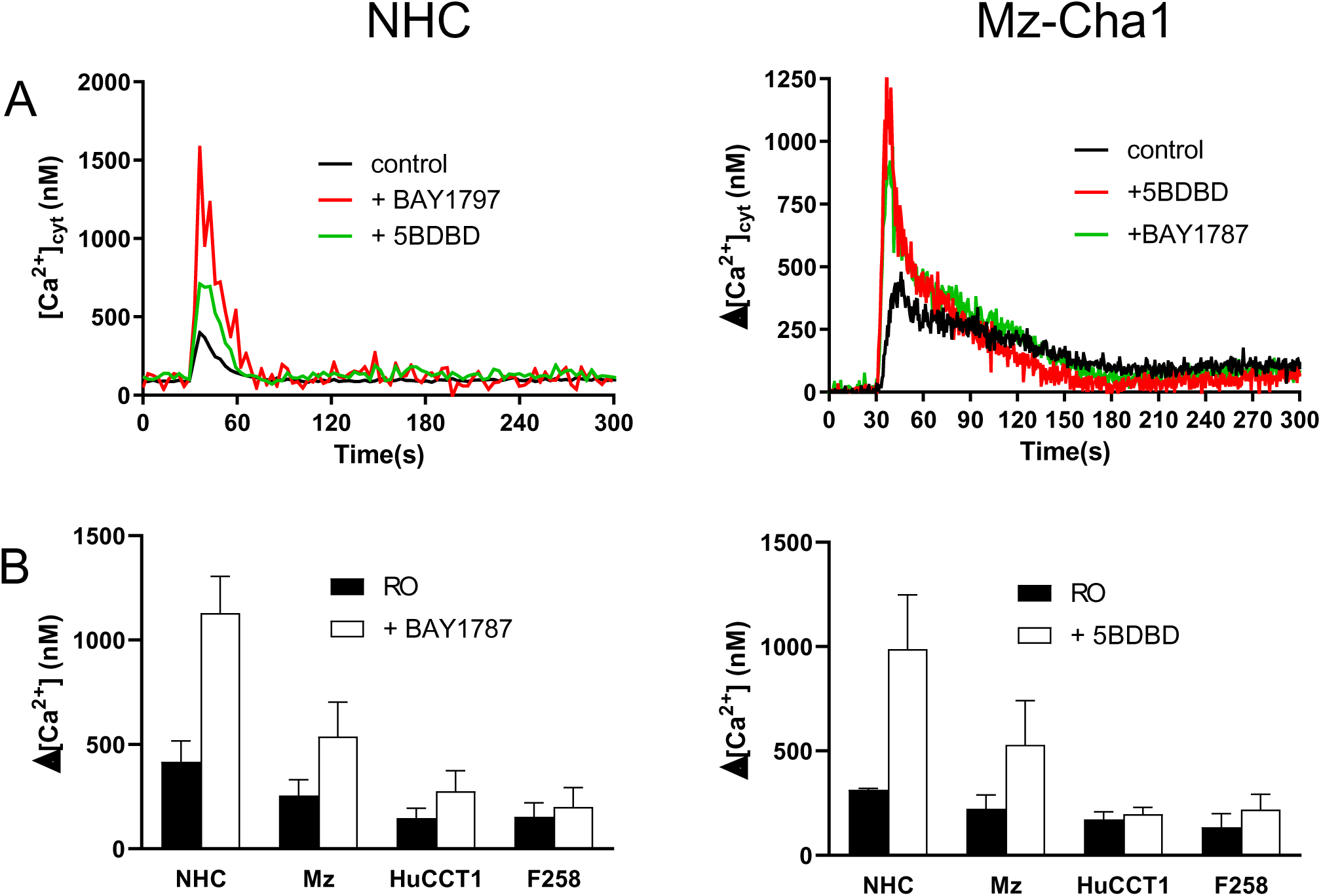
P2X4R inhibition potentiates the TGR5-induced calcium mobilisation. A. Cytosolic Calcium concentration variation (Δ[Ca^2+^]_cyt_) of NHC (left) and Mz-Cha1 cells (right), stimulated by RO5527239 (300 and 90 µM respectively) or after pretreatment by the 2 P2X4R inhibitors 5BDBD (red curves) or BAY1787 (green curves). B. Peak of RO5527239-induced calcium mobilisation in presence of BAY1787 (left) or 5BDBD (right) in the 4 cell types.

**Figure 10:**
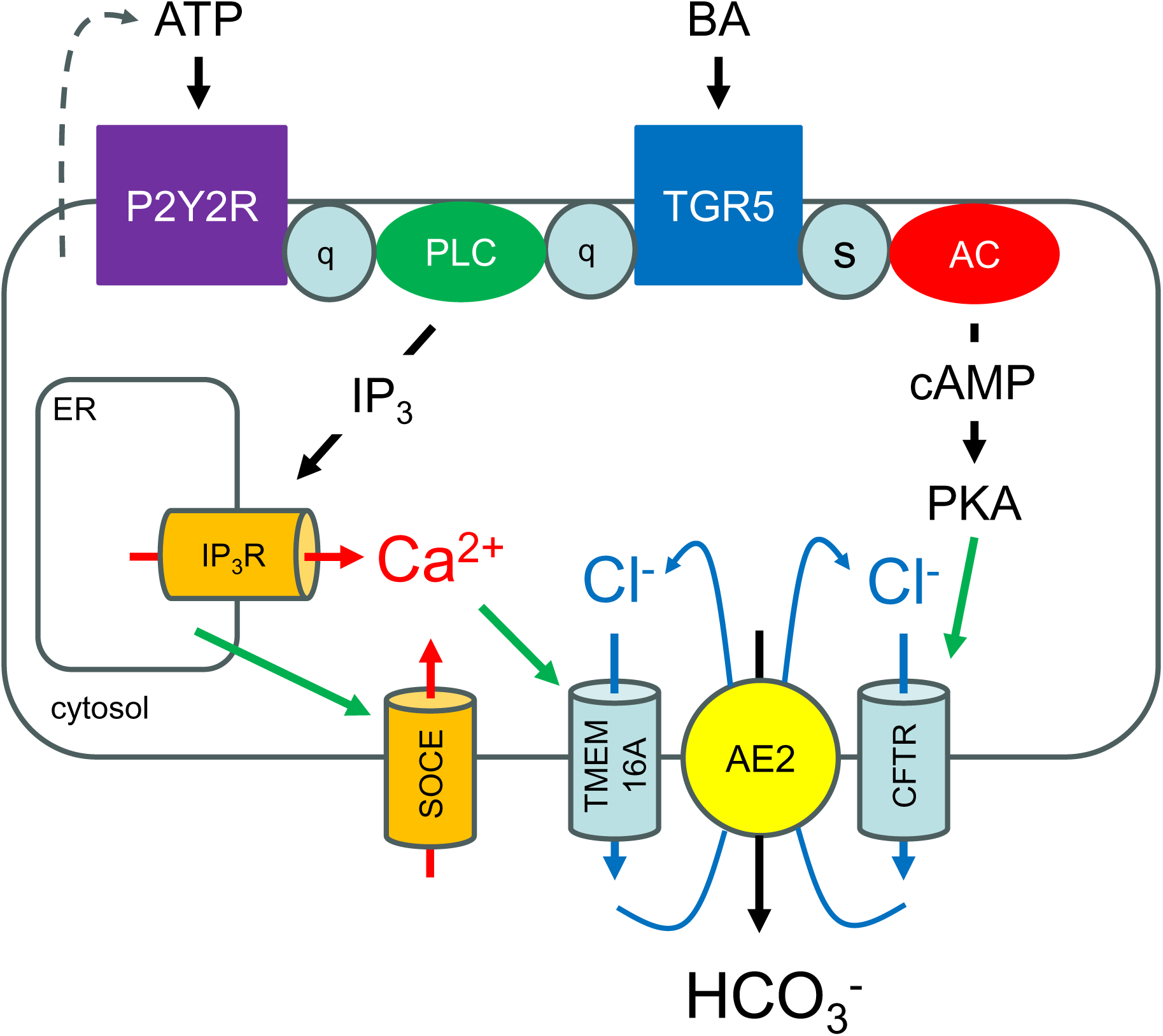
Scheme of TGR5 stimulation leading to HCO_3_^−^ secretion. Stimulation by bile acids (“BA”) of TGR5 leads to activation of two pathways depending on two G proteins: i) Gαs (“s”) allowing the activation of adenylate cyclase (“AC”), and formation of cAMP, known to activate protein kinase A (“PKA”), and thereafter of CFTR Cl^−^ channels, ii) Gαq/11 (“q”) allowing the activation of phospholipase C (“PLC”) and synthesis of IP_3_, known to induce Ca^2+^ ions release from the endoplasmic reticulum (“ER”) through IP_3_ receptors (“IP_3_R”). The emptying of the ER is known to induce the Store-Operated Calcium Entry (‘”SOCE”), resulting with the ER Ca^2+^ release to a huge increase of the cytosolic Ca^2+^ concentration, thereafter, activating the TMEM16A Cl-channel. Noteworthy, TGR5 stimulation also induces an ATP (and UTP?) secretion, leading to the reinforcement of the Gαq/11 pathways through P2Y2RR (and P2Y4RR in normal cholangiocytes) receptors. Secretion of Cl^−^ ions by CFTR and TMEM16A channels, next allows the exchange of Cl^−^ ions by bicarbonate ions through AE2. The secretion of ions is accompanied by H_2_0 secretion (not shown on the scheme). At the end, bile is more fluid and its pH is increased.

Altogether our data suggest that complex interactions between purinergic receptors and TGR5 contribute to BA-mediated Ca^2+^ responses in cholangiocytes. Thus, we showed that TGR5 stimulation induced a Ca^2+^ mobilisation by itself and an ATP (+UTP) release activating P2Y2R (and P2Y4R receptors in NHC cells), reinforcing this mobilisation.

## 4. DISCUSSION

Bile is primarily synthesized and secreted in canaliculi by hepatocytes, and then transformed during its travel along the biliary tract, in particular thanks to ion and water exchanges across the cholangiocyte epithelial barrier [44]. In the biliary tract, bile is made more fluid and alkaline, due to strong HCO_3_^−^/Cl^−^ exchanges (mediated by the AE2 protein) classically coupled with Cl^−^ exit through the CFTR channels [45]. Optimal bile fluidity and pH are not only prerequisites for bile functions in digestive processes, but also avoid the excessive accumulation of toxins and hydrophobic bile acids (BA) by flushing bile duct content, thereby protecting cholangiocytes and liver parenchyma from potential injury. The cholangiocyte compartment has also recently been considered as the site of physiological bile flow generation, instead of the hepatocyte compartment where bile would be relatively stagnant [46]. Alterations in secretory processes in bile ducts are the hallmark of cholangiopathies, including primary or secondary cholangitis a well as cystic fibrosis-associated cholangitis, during which severe cholestasis can end in the need for liver transplantation [47].

In addition to endocrine-mediated modulation [2], it is well admitted that cholangiocyte processes for bile modification occur in a BA-regulated manner. The commonly accepted view is that BA-mediated cAMP synthesis leads to CFTR activation, increasing the efflux of Cl^−^ ions (and thus also HCO_3_^−^ and H_2_O secretion). Even though this cAMP pathway was well established for several decades, it appeared that BA also induce cytosolic Ca^2+^ signals in cholangiocytes [1], and that purinergic signalling through ATP release plays a significant role in BA-induced secretory processes [48] and interferes with BA-mediated Ca^2+^ mobilisation [49, 50].

By using a TGR5 specific agonist (RO5527239), we confirmed that plasma membrane located BA-receptors induce a rise in intracellular Ca^2+^ concentration in normal human cholangiocytes and cholangiocarcinoma cell lines. For the first time, we clearly established that TGR5 was functionally coupled not only to Gαs (cAMP pathway) but also to Gαq/11 that induces an IP_3_-mediated Ca^2+^ signalling pathway. In line, specific Gαq/11 inhibition totally impaired TGR5-mediated Ca^2+^ mobilisation. Interestingly, beside cAMP-mediated activation of CFTR, Ca^2+^ signals are known to activate the Ca^2+^-activated Cl^−^ channel TMEM16A (also known as Ano1 and Anoctamin 1) [51]. Thus, Cl^−^ efflux cannot be resumed to CFTR activity, but to the co-existence of cAMP- and Ca^2+^-dependent pathways. This could be very important in the understanding of liver diseases associated with for example CF: 10-15% of CF patients develop clinical liver disease. We could hypothesize that the Ca^2+^-dependent pathway, by activating TMEM16A, might compensate, in part or totally, for the loss of the cAMP-dependent CFTR-mediated Cl^−^ secretion during CF, this remaining an open question. Interestingly, recent studies have shown that the TMEM16A Ca^2+^-activated Cl^−^ current could be 2-3 times larger than the cAMP-activated Cl^−^ current [12].

Interesting is also the role of P2X4R receptors in TGR5-mediated Ca^2+^ mobilisation. Even if the P2X4R receptor is considered as the most Ca^2+^ ion permeable channel of the P2X family, it is more a Na^+^ permeable channel, with only 11-15% of Ca^2+^ ions in the influx [52]. When using two inhibitors to block IP_3_ synthesis, ATP was no longer able to induce any Ca^2+^ rise (Fig. 49, demonstrating that P2X receptors have no role in TGR5-induced Ca^2+^ mobilisation. However, when we pre-treated cells with two specific P2X4R inhibitors, TGR5-mediated Ca^2+^ mobilisation was increased (Fig 9) This suggests that P2X4R receptors are present and active in cholangiocytes, but presumably not as a Ca^2+^ channel and rather as a modulator of Ca^2+^ mobilisation. Noteworthy, the genetically close P2X7 channel, known also to transport Na^+^ and Ca^2+^ has been shown to induce the depolarization of B DT40 cells plasma membrane [53]. Thus, we could speculate that by decreasing the membrane potential, P2X4R receptor activation would decrease the Ca^2+^ driving force and reduce the SOCE.

TGR5-mediated functions in cholangiocytes, in addition to Cl^−^ secretion, also include the modulation of cell proliferation and apoptosis [23]. Interestingly, TGR5-induced stimulation of cholangiocyte proliferation turned to be cAMP-independent [23], opening possible involvement of Ca^2+^ signalling. Indeed, metallo-protease and kinase activation, as well as ROS generation, all of these being described as triggered by TGR5 stimulation, have all been reported as Ca^2+^-dependent. As regard to the resting [Ca^2+^]_cyt._, we found that cholangiocarcinoma cell lines had a significantly decreased resting [Ca^2+^]_cyt_, but also a weaker Store-Operated Calcium Entry mainly due to Orai1. Although altered calcium signalling has been reported as a hallmark of cancer cells [54], it remains undefined if the observed differences play a role in the carcinogenesis process or would be secondary to cell transformation. Interestingly, the downregulation of Orai1, inducing a decreased-SOCE, decreased apoptosis in human prostate cancer cells [55]. Direct impact of TGR5-elicited Ca^2+^ signalling on cholangiocyte proliferation, either in regenerative or neoplasic contexts, however, still remains to be explored.

Although crosstalk between BA, purinergic and calcium signalling in the context of cholangiocyte secretion had already been reported [9, 48], our study deciphers in details such a crosstalk involving after BA challenge, the BA receptor TGR5 coupled to Gαq11, ATP release, P2Y2R (and P2Y4R in normal cholangiocytes) engagement, resulting in both calcium release from the ER and calcium entry (resumed in cartoon of Fig 10). On the top of this pathway, we highlighted further complex feedback control through the ATP-stimulated P2X4R receptor. Specific P2Y2R inhibition only partly blunted TGR5-dependent Ca^2+^ mobilisation, indicating that both receptors contribute on their own to BA-induced calcium mobilization. Nevertheless, the 2 pathways crosstalk through ATP release by cholangiocytes. As ATP secretion has been reported to be CFTR-dependent [48], a simple scheme could be that after TGR5 stimulation, Gαq/11 activation leads to IP_3_ synthesis and Ca^2+^ release from the ER, resulting in the opening of Store-Operated Calcium channels. This would allow the start of a cytosolic Ca^2+^ concentration increase and the opening of TMEM16A channels. In parallel, TGR5-induced Gαs activation and cAMP production would next favor CFTR channels opening that elicit Cl^−^ ions efflux and ATP release. This ATP will activate P2Y2R and P2Y4R receptors, reinforcing IP_3_ synthesis and downstream Ca^2+^ pathways. Noteworthy, the use of P2Y2R activator (denufosol) to activate the TMEM16A-dependent Cl^−^ secretion in CF has been documented [56]. Thus, denufosol was able to improve the hydration of the pulmonary airways and the cilia beat frequency by stimulating P2Y2R receptor and next activation of TMEM16A.

Notheworthy in normal human cholangiocytes, UTP induced a Ca^2+^ mobilisation through P2Y2R and P2Y4R receptors, when only P2Y2R are activated in cell lines. Further experiments are needed to understand if the loss of the P2Y4R receptors could be associated to cholangiocarcinoma genesis or would be a consequence of this neoplasic transformation.

## 5. CONCLUSIONS

By sig normal cholangiocytes and cholangiocarcinoma cell lines, we showed that beside the well-documented cAMP pathway induced by TGR5-dependent BA challenge in cholangiocytes, a Ca^2+^ signalling pathway is also coupled with this receptor. Our study put forward a complex crosstalk between TGR5-dependent and purinergic input converging on calcium signals, that should be considered in pathophysiological contexts for possible targeting of secretory or proliferation processes in cholangiocytes. Indeed, this alternative Ca^2+^ pathways could activate a Cl^−^ secretion that might be able to compensate for the loss of CFTR channel activity in CF.

## ABBREVIATIONS

TGR5: Takeda G-protein Receptor 5
CFTR: Cystic Fibrosis Transmembrane conductance Regulator
ATP: Adenosine Trisphosphate
UTP: Uridine Trisphosphate
NTP: Nucleotide Trisphosphate
cAMP: cyclic Adenosine Monophosphate
IP_3_: inositol 1,4,5-triphosphate
GPCR: G-Protein Coupled Receptor
BA: Bile Acid
SOCE: Store-Operated Calcium Entry
NHC: Normal Human Cholangiocytes
CCA: Cholangiocarcinoma
ER: endoplasmic reticulum
[Ca^2+^]_cyt_: cytosolic calcium concentration
SERCA: sarco/endoplasmic reticulum Ca^2+^ ATPase
TG: thapsigargin
PL: Phospholipase
AE2: Anion Exchange Protein 2
P2Y2R: P2Y2 receptor

## CRediT authorship contribution statement

**CHEN Xianmeng** : investigation, formal analysis. **AL SHEBEL Amr**: investigation, formal analysis. **PEBRIER Thibault**: investigation, formal analysis. **TORDJMANN Thierry**: funding acquisition, writing – review & editing. **DELLIS Olivier**: conceptualization, methodology, writing – original draft, writing - review & editing, supervision, project administration, visualization.

## AKNOWLEDGMENTS

This study was funded by Institut National de la Santé et de la Recherche Médicale, Université Paris-Saclay and Agence Nationale de la Recherche (Grant Number 15-CE14-0007-01)

## REFERENCES

[1] M.F. Leite, M.T. Guera, V.A. Andrade, M.H. Nathanson, Signaling Pathways in Biliary Epithelial Cells, 2015.

[2] G. Alpini, C.D. Ulrich, J.O. Phillips, L.D. Pham, L.J. Miller, N.F. Larusso, Up-Regulation of Secretin Receptor Gene-Expression in Rat Cholangiocytes after Bile-Duct Ligation, Am J Physiol, 266 (1994) G922–G928.

[3] R. Lenzen, G. Alpini, N. Tavoloni, Secretin Stimulates Bile Ductular Secretory Activity through the Camp System, Am J Physiol, 263 (1992) G527–G532.

[4] J.M. McGill, S. Basavappa, T.W. Gettys, J.G. Fitz, Secretin activates Cl-channels in bile duct epithelial cells through a cAMP-dependent mechanism, Am J Physiol, 266 (1994) G731–736.

[5] D. Alvaro, W.K. Cho, A. Mennone, J.L. Boyer, Effect of secretion on intracellular pH regulation in isolated rat bile duct epithelial cells, J Clin Invest, 92 (1993) 1314–1325.

[6] A. Mennone, D. Alvaro, W. Cho, J.L. Boyer, Isolation of small polarized bile duct units, Proc Natl Acad Sci U S A, 92 (1995) 6527–6531.

[7] S.K. Roberts, S.M. Kuntz, G.J. Gores, N.F. LaRusso, Regulation of bicarbonate-dependent ductular bile secretion assessed by lumenal micropuncture of isolated rodent intrahepatic bile ducts, Proc Natl Acad Sci U S A, 90 (1993) 9080–9084.

[8] T. Flass, M.R. Narkewicz, Cirrhosis and other liver disease in cystic fibrosis, J Cyst Fibros, 12 (2013) 116–124.

[9] J.A. Dranoff, A.I. Masyuk, E.A. Kruglov, N.F. LaRusso, M.H. Nathanson, Polarized expression and function of P2Y ATP receptors in rat bile duct epithelia, Am J Physiol Gastrointest Liver Physiol, 281 (2001) G1059–1067.

[10] R.S. Chari, S.M. Schutz, J.E. Haebig, G.H. Shimokura, P.B. Cotton, J.G. Fitz, W.C. Meyers, Adenosine nucleotides in bile, Am J Physiol, 270 (1996) G246–252.

[11] M.H. Nathanson, A.D. Burgstahler, A. Masyuk, N.F. Larusso, Stimulation of ATP secretion in the liver by therapeutic bile acids, Biochem J, 358 (2001) 1–5.

[12] A.K. Dutta, A.K. Khimji, C. Kresge, A. Bugde, M. Dougherty, V. Esser, Y. Ueno, S.S. Glaser, G. Alpini, D.C. Rockey, A.P. Feranchak, Identification and functional characterization of TMEM16A, a Ca2+-activated Cl-channel activated by extracellular nucleotides, in biliary epithelium, J Biol Chem, 286 (2011) 766–776.

[13] Q. Li, A. Dutta, C. Kresge, A. Bugde, A.P. Feranchak, Bile acids stimulate cholangiocyte fluid secretion by activation of transmembrane member 16A Cl(-) channels, Hepatology, 68 (2018) 187–199.

[14] G.H. Shimokura, J.M. McGill, T. Schlenker, J.G. Fitz, Ursodeoxycholate increases cytosolic calcium concentration and activates Cl-currents in a biliary cell line, Gastroenterology, 109 (1995) 965–972.

[15] Y. Kawamata, R. Fujii, M. Hosoya, M. Harada, H. Yoshida, M. Miwa, S. Fukusumi, Y. Habata, T. Itoh, Y. Shintani, S. Hinuma, Y. Fujisawa, M. Fujino, A G protein-coupled receptor responsive to bile acids, J Biol Chem, 278 (2003) 9435–9440.

[16] V. Keitel, K. Cupisti, C. Ullmer, W.T. Knoefel, R. Kubitz, D. Haussinger, The membrane-bound bile acid receptor TGR5 is localized in the epithelium of human gallbladders, Hepatology, 50 (2009) 861–870.

[17] C. Guo, W.D. Chen, Y.D. Wang, TGR5, Not Only a Metabolic Regulator, Front Physiol, 7 (2016) 646.

[18] M. Watanabe, S.M. Houten, C. Mataki, M.A. Christoffolete, B.W. Kim, H. Sato, N. Messaddeq, J.W. Harney, O. Ezaki, T. Kodama, K. Schoonjans, A.C. Bianco, J. Auwerx, Bile acids induce energy expenditure by promoting intracellular thyroid hormone activation, Nature, 439 (2006) 484–489.

[19] A. Perino, H. Demagny, L. Velazquez-Villegas, K. Schoonjans, Molecular Physiology of Bile Acid Signaling in Health, Disease, and Aging, Physiol Rev, 101 (2021) 683–731.

[20] N. Péan, I. Doignon, I. Garcin, A. Besnard, B. Julien, B. Liu, S. Branchereau, A. Spraul, C. Guettier, L. Humbert, K. Schoonjans, D. Rainteau, T. Tordjmann, The receptor TGR5 protects the liver from bile acid overload during liver regeneration in mice, Hepatology, 58 (2013) 1451–1460.

[21] G. Merlen, V. Bidault-Jourdainne, N. Kahale, M. Glenisson, J. Ursic-Bedoya, I. Doignon, I. Garcin, L. Humbert, D. Rainteau, T. Tordjmann, Hepatoprotective impact of the bile acid receptor TGR5, Liver Int, 40 (2020) 1005–1015.

[22] T. Maruyama, Y. Miyamoto, T. Nakamura, Y. Tamai, H. Okada, E. Sugiyama, H. Itadani, K. Tanaka, Identification of membrane-type receptor for bile acids (M-BAR), Biochem Biophys Res Commun, 298 (2002) 714–719.

[23] M. Reich, K. Deutschmann, A. Sommerfeld, C. Klindt, S. Kluge, R. Kubitz, C. Ullmer, W.T. Knoefel, D. Herebian, E. Mayatepek, D. Haussinger, V. Keitel, TGR5 is essential for bile acid-dependent cholangiocyte proliferation in vivo and in vitro, Gut, 65 (2016) 487–501.

[24] K. Deutschmann, M. Reich, C. Klindt, C. Droge, L. Spomer, D. Haussinger, V. Keitel, Bile acid receptors in the biliary tree: TGR5 in physiology and disease, Biochim Biophys Acta Mol Basis Dis, 1864 (2018) 1319–1325.

[25] V. Keitel, M. Reich, D. Haussinger, TGR5: pathogenetic role and/or therapeutic target in fibrosing cholangitis?, Clin Rev Allergy Immunol, 48 (2015) 218–225.

[26] V. Bidault-Jourdainne, G. Merlen, M. Glénisson, I. Doignon, I. Garcin, N. Péan, R. Boisgard, J. Ursic-Bedoya, M. Serino, C. Ullmer, L. Humbert, A. Abdelrafee, N. Golse, E. Vibert, J.C. Duclos-Vallée, D. Rainteau, T. Tordjmann, TGR5 controls bile acid composition and gallbladder function to protect the liver from bile acid overload, JHEP Rep, 3 (2021) 100214.

[27] M. Reich, L. Spomer, C. Klindt, K. Fuchs, J. Stindt, K. Deutschmann, J. Höhne, E. Liaskou, J.R. Hov, T.H. Karlsen, U. Beuers, J. Verheij, S. Ferreira-Gonzalez, G. Hirschfield, S.J. Forbes, C. Schramm, I. Esposito, D. Nierhoff, P. Fickert, C.D. Fuchs, M. Trauner, M. García-Beccaria, G. Gabernet, S. Nahnsen, J.P. Mallm, M. Vogel, K. Schoonjans, T. Lautwein, K. Köhrer, D. Häussinger, T. Luedde, M. Heikenwalder, V. Keitel, Downregulation of TGR5 (GPBAR1) in biliary epithelial cells contributes to the pathogenesis of sclerosing cholangitis, J Hepatol, 75 (2021) 634–646.

[28] H. Dehmlow, R. Alvarez Sánchez, S. Bachmann, C. Bissantz, F. Bliss, K. Conde-Knape, M. Graf, R.E. Martin, U. Obst Sander, S. Raab, H.G. Richter, S. Sewing, U. Sprecher, C. Ullmer, P. Mattei, Discovery and optimisation of 1-hydroxyimino-3,3-diphenylpropanes, a new class of orally active GPBAR1 (TGR5) agonists, Bioorg Med Chem Lett, 23 (2013) 4627–4632.

[29] G. Grynkiewicz, M. Poenie, R.Y. Tsien, A new generation of Ca2+ indicators with greatly improved fluorescence properties, J Biol Chem, 260 (1985) 3440–3450.

[30] A. Djillani, O. Nusse, O. Dellis, Characterization of novel store-operated calcium entry effectors, Biochim Biophys Acta, 1843 (2014) 2341–2347.

[31] E. Morel-Chany, C. Guillouzo, G. Trincal, M.F. Szajnert, “Spontaneous” neoplastic transformation in vitro of epithelial cell strains of rat liver: cytology, growth and enzymatic activities, Eur J Cancer, 14 (1978) 1341–1352.

[32] O. Erice, I. Labiano, A. Arbelaiz, A. Santos-Laso, P. Munoz-Garrido, R. Jimenez-Aguero, P. Olaizola, A. Caro-Maldonado, N. Martin-Martin, A. Carracedo, E. Lozano, J.J. Marin, C.J. O’Rourke, J.B. Andersen, J. Llop, V. Gomez-Vallejo, D. Padro, A. Martin, M. Marzioni, L. Adorini, M. Trauner, L. Bujanda, M.J. Perugorria, J.M. Banales, Differential effects of FXR or TGR5 activation in cholangiocarcinoma progression, Biochim Biophys Acta Mol Basis Dis, 1864 (2018) 1335–1344.

[33] Y. Wang, H. Aoki, J. Yang, K. Peng, R. Liu, X. Li, X. Qiang, L. Sun, E.C. Gurley, G. Lai, L. Zhang, G. Liang, M. Nagahashi, K. Takabe, W.M. Pandak, P.B. Hylemon, H. Zhou, The role of sphingosine 1-phosphate receptor 2 in bile-acid-induced cholangiocyte proliferation and cholestasis-induced liver injury in mice, Hepatology, 65 (2017) 2005–2018.

[34] J.W. Putney, Jr., A model for receptor-regulated calcium entry, Cell Calcium, 7 (1986) 1–12.

[35] J.I. Bruce, D.I. Yule, T.J. Shuttleworth, Ca2+-dependent protein kinase--a modulation of the plasma membrane Ca2+-ATPase in parotid acinar cells, J Biol Chem, 277 (2002) 48172–48181.

[36] E.R. Lazarowski, J.I. Sesma, L. Seminario-Vidal, S.M. Kreda, Molecular mechanisms of purine and pyrimidine nucleotide release, Adv Pharmacol, 61 (2011) 221–261.

[37] J.R. Goree, E.G. Lavoie, M. Fausther, J.A. Dranoff, Expression of mediators of purinergic signaling in human liver cell lines, Purinergic Signal, 10 (2014) 631–638.

[38] C. Elsing, T. Georgiev, C.A. Hubner, R. Boger, W. Stremmel, T. Schlenker, Extracellular ATP induces cytoplasmic and nuclear Ca2+ transients via P2Y2 receptor in human biliary epithelial cancer cells (Mz-Cha-1), Anticancer Res, 32 (2012) 3759–3767.

[39] S. Dolovcak, S.L. Waldrop, J.G. Fitz, G. Kilic, 5-Nitro-2-(3-phenylpropylamino)benzoic acid (NPPB) stimulates cellular ATP release through exocytosis of ATP-enriched vesicles, J Biol Chem, 284 (2009) 33894–33903.

[40] M. Fausther, E. Gonzales, J.A. Dranoff, Role of purinergic P2X receptors in the control of liver homeostasis, Wiley Interdiscip Rev Membr Transp Signal, 1 (2012) 341–348.

[41] J.H. Tabibian, A.I. Masyuk, T.V. Masyuk, S.P. O’Hara, N.F. LaRusso, Physiology of cholangiocytes, Compr Physiol, 3 (2013) 541–565.

[42] I. von Kugelgen, Pharmacological profiles of cloned mammalian P2Y-receptor subtypes, Pharmacol Ther, 110 (2006) 415–432.

[43] R.B. Doctor, T. Matzakos, R. McWilliams, S. Johnson, A.P. Feranchak, J.G. Fitz, Purinergic regulation of cholangiocyte secretion: identification of a novel role for P2X receptors, Am J Physiol Gastrointest Liver Physiol, 288 (2005) G779–786.

[44] J.L. Boyer Bile formation and secretion, Wiley 2013.

[45] J.M. Banales, J. Prieto, J.F. Medina, Cholangiocyte anion exchange and biliary bicarbonate excretion, World J Gastroenterol, 12 (2006) 3496–3511.

[46] N. Vartak, G. Guenther, F. Joly, A. Damle-Vartak, G. Wibbelt, J. Fickel, S. Jörs, B. Begher-Tibbe, A. Friebel, K. Wansing, A. Ghallab, M. Rosselin, N. Boissier, I. Vignon-Clementel, C. Hedberg, F. Geisler, H. Hofer, P. Jansen, S. Hoehme, D. Drasdo, J.G. Hengstler, Intravital Dynamic and Correlative Imaging of Mouse Livers Reveals Diffusion-Dominated Canalicular and Flow-Augmented Ductular Bile Flux, Hepatology, 73 (2021) 1531–1550.

[47] J.M. Banales, J.J.G. Marin, A. Lamarca, P.M. Rodrigues, S.A. Khan, L.R. Roberts, V. Cardinale, G. Carpino, J.B. Andersen, C. Braconi, D.F. Calvisi, M.J. Perugorria, L. Fabris, L. Boulter, R.I.R. Macias, E. Gaudio, D. Alvaro, S.A. Gradilone, M. Strazzabosco, M. Marzioni, C. Coulouarn, L. Fouassier, C. Raggi, P. Invernizzi, J.C. Mertens, A. Moncsek, S. Rizvi, J. Heimbach, B.G. Koerkamp, J. Bruix, A. Forner, J. Bridgewater, J.W. Valle, G.J. Gores, Cholangiocarcinoma 2020: the next horizon in mechanisms and management, Nat Rev Gastroenterol Hepatol, 17 (2020) 557–588.

[48] R. Fiorotto, C. Spirli, L. Fabris, M. Cadamuro, L. Okolicsanyi, M. Strazzabosco, Ursodeoxycholic acid stimulates cholangiocyte fluid secretion in mice via CFTR-dependent ATP secretion, Gastroenterology, 133 (2007) 1603–1613.

[49] J.M. Kowal, K.A. Haanes, N.M. Christensen, I. Novak, Bile acid effects are mediated by ATP release and purinergic signalling in exocrine pancreatic cells, Cell Commun Signal, 13 (2015) 28.

[50] Q. Li, A. Dutta, C. Kresge, A. Bugde, A.P. Feranchak, Bile acids stimulate cholangiocyte fluid secretion by activation of transmembrane member 16A Cl, Hepatology, 68 (2018) 187–199.

[51] Y. Jang, U. Oh, Anoctamin 1 in secretory epithelia, Cell Calcium, 55 (2014) 355–361.

[52] L. Stokes, J.A. Layhadi, L. Bibic, K. Dhuna, S.J. Fountain, P2X4 Receptor Function in the Nervous System and Current Breakthroughs in Pharmacology, Front Pharmacol, 8 (2017) 291.

[53] M. Tsukimoto, H. Harada, A. Ikari, K. Takagi, Involvement of chloride in apoptotic cell death induced by activation of ATP-sensitive P2X7 purinoceptor, J Biol Chem, 280 (2005) 2653–2658.

[54] S. Marchi, P. Pinton, Alterations of calcium homeostasis in cancer cells, Curr Opin Pharmacol, 29 (2016) 1–6.

[55] M. Flourakis, V. Lehen’kyi, B. Beck, M. Raphael, M. Vandenberghe, F.V. Abeele, M. Roudbaraki, G. Lepage, B. Mauroy, C. Romanin, Y. Shuba, R. Skryma, N. Prevarskaya, Orai1 contributes to the establishment of an apoptosis-resistant phenotype in prostate cancer cells, Cell Death Dis, 1 (2010) e75.

[56] H.L. Danahay, S. Lilley, R. Fox, H. Charlton, J. Sabater, B. Button, C. McCarthy, S.P. Collingwood, M. Gosling, TMEM16A Potentiation: A Novel Therapeutic Approach for the Treatment of Cystic Fibrosis, Am J Respir Crit Care Med, 201 (2020) 946–954.

